# Multi-omics data analysis implicating epigenetic inheritance in evolution and disease

**DOI:** 10.1101/770099

**Authors:** Abhay Sharma

## Abstract

Recent evidence surprisingly suggests existence of germline mediated epigenetic inheritance in diverse species including mammals. The evolutionary and health implications as well as the mechanistic plausibility of epigenetic inheritance are subjects of immense current interest and controversy, with integrative analysis expected to provide valuable insights. Here, an unbiased gene set enrichment analysis of existing multi-omics data is presented that readily supports a role of sperm DNA methylome in evolution and disease, as also in developmental mechanisms. In mice, differentially methylated sperm genes in cold induced inheritance specifically overrepresent genes associated with cold adaptation. Similarly, in humans, differentially methylated sperm genes associate with disease and adaptation in general, with specific disease association supported by prior evidence. Further, the sperm genes, like disease and adaptation genes, overrepresent genes known to exhibit higher mutability, loss-of-function intolerance, and haploinsufficiency. Finally, both mouse and human sperm genes show enrichment for genes that retain sperm methylation during development and are developmentally expressed. Together, the present analysis provides one-stop evidence to suggest that sperm DNA methylome acts as a melting pot of gene-environment interaction, inheritance, evolution, and health and disease.

## Introduction

Of immense current interest and debate, recent evidence suggests that parental environmental exposure may induce phenotypic effects in unexposed progeny through germline epigenetic inheritance in diverse species including mammals^1,2^. Sperm DNA methylation, known to partially survive developmental demethylation, is considered as one of the epigenetic factors that may mediate inheritance by influencing development^1,3,4^. Also, epigenetic inheritance holds the potential to impact evolution and disease^5-7^. With DNA methylation known to influence DNA sequence mutability, environmentally induced inheritance may possibly lead to genetic fixation of traits^5,6,8^. However, the evolutionary and biomedical significance, and developmental mechanisms of epigenetic inheritance remain overall highly controversial^1,9-11^. Integrative analysis of diverse data may offer key to the much needed understanding^1,5^. Here, I show that sperm DNA methylome centred gene set analysis of existing multi-omics data does provide the anticipated insights. In mice, differentially methylated sperm genes in cold induced inheritance^12^ specifically overrepresent genes associated with latitudinal adaptation^13,14^. In humans, differentially methylated sperm genes in recreational cannabis users and in fathers of autistic children, consistent with previous evidence^15,16^, specifically overrepresent genes associated with autism and/or schizophrenia. Remarkably, the sperm genes in human under various conditions combined show enrichment of disease and adaptation associated genes in general. Further, these genes, like human disease associated genes combined, also overrepresent genes exhibiting high mutability, loss-of-function intolerance, and haploinsufficiency. Lastly, differentially methylated sperm genes in various mouse models of inheritance or human conditions combined show enrichment for genes that retain sperm methylation during development or are developmentally expressed.

## Results and discussion

Evolutionary and disease implications of epigenetic inheritance were examined upfront. For evolutionary significance, I tested the overlap between genes showing differential methylation in sperm in various mouse models of paternal inheritance and genes associated with evolutionary adaptation in mice. The environmental factors used to trigger inheritance in these models included low-protein diet^17^ (LPD), cold exposure^12^ (CE), and a combination of high-fat diet and the diabetes phenotype inducing compound streptozotocin^18^ (H+S), with the studies identified from literature-wide search using the criteria of demonstrated physiological and gene regulatory effects in the offspring. The evolutionarily related genes comprised of that showing single nucleotide polymorphism (SNP) association and allele specific expression (ASE), as also differentially expressed genes (DEG), in latitudinal adaptation^13,14^. I tested the null hypothesis that the magnitude of overlap between cold adaptation genes and LPD, CE or H+S is the same as that between the former and sperm genes in all three models combined. Alternatively, the magnitude of overlap with CE alone is higher, supporting impact of CE in latitudinal adaptation. Remarkably, CE, not LPD or H+S, was found to overrepresent adaptation genes (**Fig. 1a**). Though the overrepresentation remained significant for only ASE and DEG, not SNP, following *p* value correction, the CE enrichment of latitudinal genes combined did survive multiple testing (**Fig. 1a**). Together, the analysis supported a role of epigenetic inheritance in evolutionary adaptation.

**Fig. 1.**
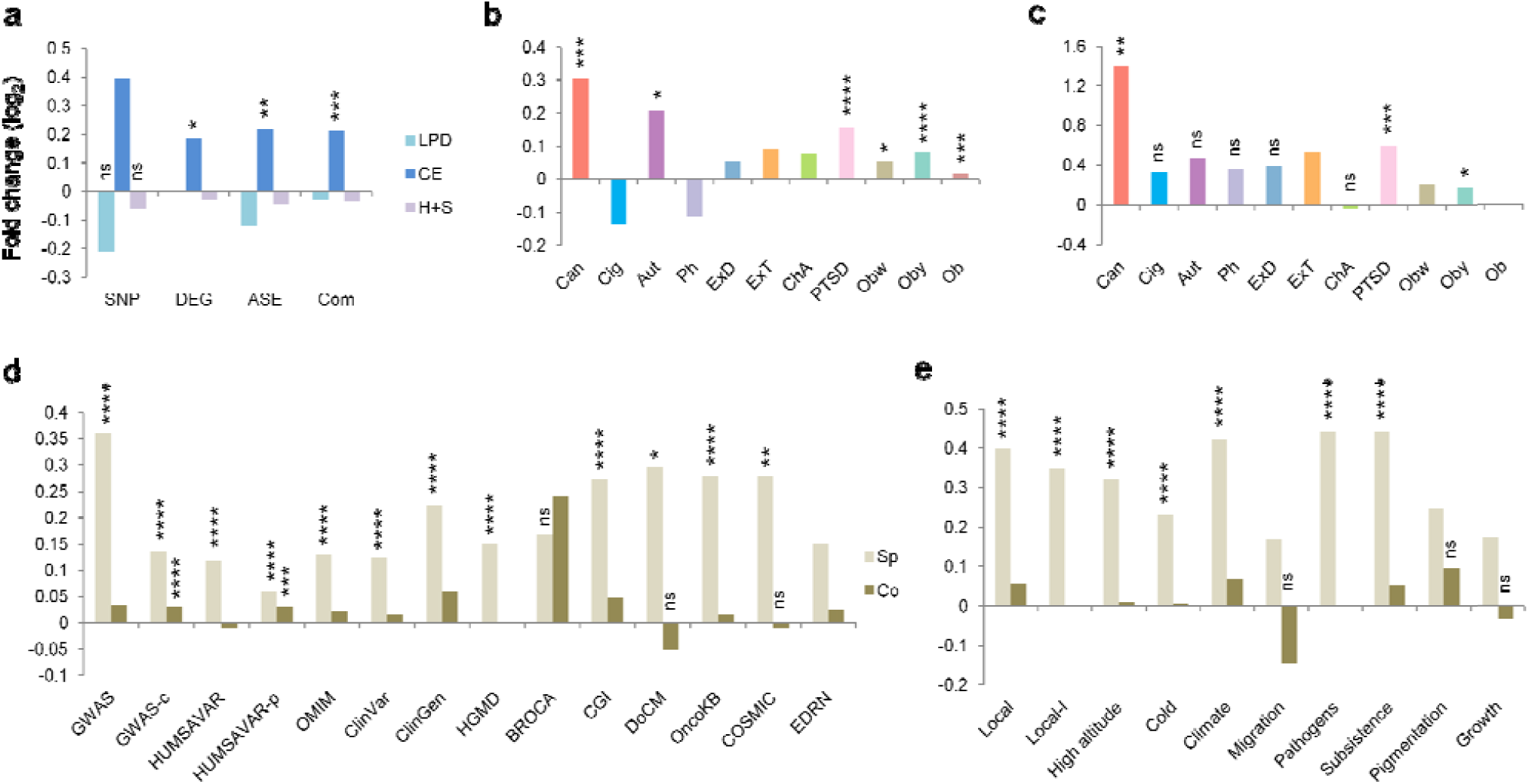
Overlap of sperm gene sets with adaptation and disease associated genes. Fold enrichment or depletion of individual mouse model sperm genes in latitudinal adaptation (**a)**, human sperm genes under each condition in ASD (**b**), or SCZ (**c**), and human sperm genes under all conditions combined in disease databases (**d**) or individual adaptations (**e**). Co, EWAS control; Com, combined set of SNP, ASE, and DEG genes; ns, unadjusted *p* not significant; Sp, all 11 human sperm conditions (b, c) combined. *, **, ***, and **** indicate figure panel-wide Bonferroni-adjusted *p* <0.05, <0.01, <0.001, and <0.0001, in that order. No mark above bars indicate unadjusted *p* <0.05. Note unique enrichment of CE across cold adaptation gene sets (a). Also, appreciate enrichment of Can both in ASD (b) and SCZ (c), and of Aut in ASD (b). Besides, note enrichment of Sp, not Co, across disease (d) and adaptation (e) genes. Mouse models (a) and human conditions (b, c) are arranged on the basis of, from left to right, increasing number of total sperm genes. Sp and Co (d, e) contained equal number of genes. Gene list sources, along with remarks, if any, and the overlapping genes found are presented in **Supplementary Table 1**. Other abbreviations and details as mentioned in the text.

For disease significance, I first determined the overlap between genes showing differential methylation in sperm under various conditions and genes associated with certain diseases implicated earlier. The sperm samples included that of recreational users of cannabis^19^ (Can), cigarette smokers^20^ (Cig), enriched-risk cohort of fathers of autistic children^15^ (Aut), men undergoing fertility treatment and showing association between differential sperm methylation and preconception urinary phthalate concentration^21^ (Ph), exercise trained (ExT) and detrained (ExD) individuals^22^, men with experience of childhood abuse^23^ (ChA), veterans with posttraumatic stress disorder^24^ (PTSD), and obese men before (Ob) and after one week (Obw) or one year (Oby) of bariatric surgery^25^. The corresponding studies were identified through manual search for relevant literature. Prior evidence suggested that sperm methylation in Can and Aut relate to genes associated with both autism spectrum disorder (ASD) and schizophrenia (SCZ), and with ASD, in that order^15,16^. I thus compared the extent of overlap between database genes representing germline *de novo* variants in ASD or SCZ and sperm genes under each of the 11 conditions mentioned above with that between the former and sperm genes under all the conditions combined. Interestingly, both Can and Aut were found to overrepresent ASD (**Fig. 1b**), with Can also showing enrichment of SCZ (**Fig. 1c**). Besides, it was notable that both PTSD and obesity associated sperm genes, Ob, Obw, and/ or Oby, overrepresented ASD (**Fig. 1b**) and SCZ (**Fig. 1c**). This is consistent with the original findings that PTSD sperm genes associate with offspring mental health diagnosis^24^, and obesity associated sperm genes overrepresent^25^ the very same Aut genes used in the present analysis.

The disease-specific association of sperm genes above (**Fig. 1b, c**) led to the speculation that differential methylation in sperm signifies a role of epigenetics in disease and adaptation in general. I therefore tested the null hypothesis that the degree of overlap between differential sperm genes in human subjects under all the 11 conditions combined and human disease database genes or adaptation associated genes is the same as that between the former and genes in the whole human genome. The alternative hypothesis was that the extent of overlap with disease and adaptation is greater, supporting the prediction. To control for gene methylation bias, the test was repeated by replacing differential sperm genes with same number of randomly selected epigenome-wide association study (EWAS) genes showing altered DNA methylation in disease and traits. Remarkably, the sperm gene set, not the EWAS control, showed enrichment of genes associated with disease (**Fig. 1d**) and adaptation (**Fig. 1e**), in general. The robustness of disease association (**Fig. 1d**) was confirmed by the observation that, as expected, the enrichment fold change was higher for a hand curated set of stronger genome-wide association study genes (GWAS) than for the total NHGRI-EBI catalog that include both stronger as well as weaker, sub-genome-wide significant signals (GWAS-c). Similarly, the fold change was higher for missense variants reported to be implicated in disease (HUMSAVAR), compared to missense variants not reported to be implicated in disease (HUMSAVAR-p). In adaptation analysis (**Fig. 1e**), the robustness of the results was reassured by the finding that, as expected, the enrichment fold change was greater for genes showing higher strength association in local adaptation (Local) than for lower strength genes (Local-l).

Disease associated genes have previously been associated with high mutability, loss-of-function intolerance, and haploinsufficiency^26-28^. To further examine disease significance of sperm methylome in humans, I tested the overlap between combined set of human sperm genes or disease associated genes with gene sets representing different genome-wide estimates of mutability, loss-of-function intolerance, and haploinsufficiency. As above (**Fig. 1d, e**), EWAS control was used for sperm analysis. Similarly, for disease analysis, the two previous pairs (**Fig. 1d**) GWAS and GWAS-c, and HUMSAVAR and HUMSAVAR-p were used. Also, like above (**Fig. 1d, e**), genes in the whole human genome was used as background population. Remarkably, it was found that both sperm and disease gene sets, not the respective controls, are enriched for genes representing higher mutability (**Fig. 2a-c**), loss-of-function intolerance (**Fig. 2d-f**), and haploinsufficiency (**Fig. 2g-i**), in general. These findings suggested that genes associated with evolutionary adaptation may also exhibit similar mutational profiles. I tested this using the previous (**Fig. 1e**) gene set pair Local and Local-l, and found supporting evidence (**Fig. 3**).

**Fig. 2.**
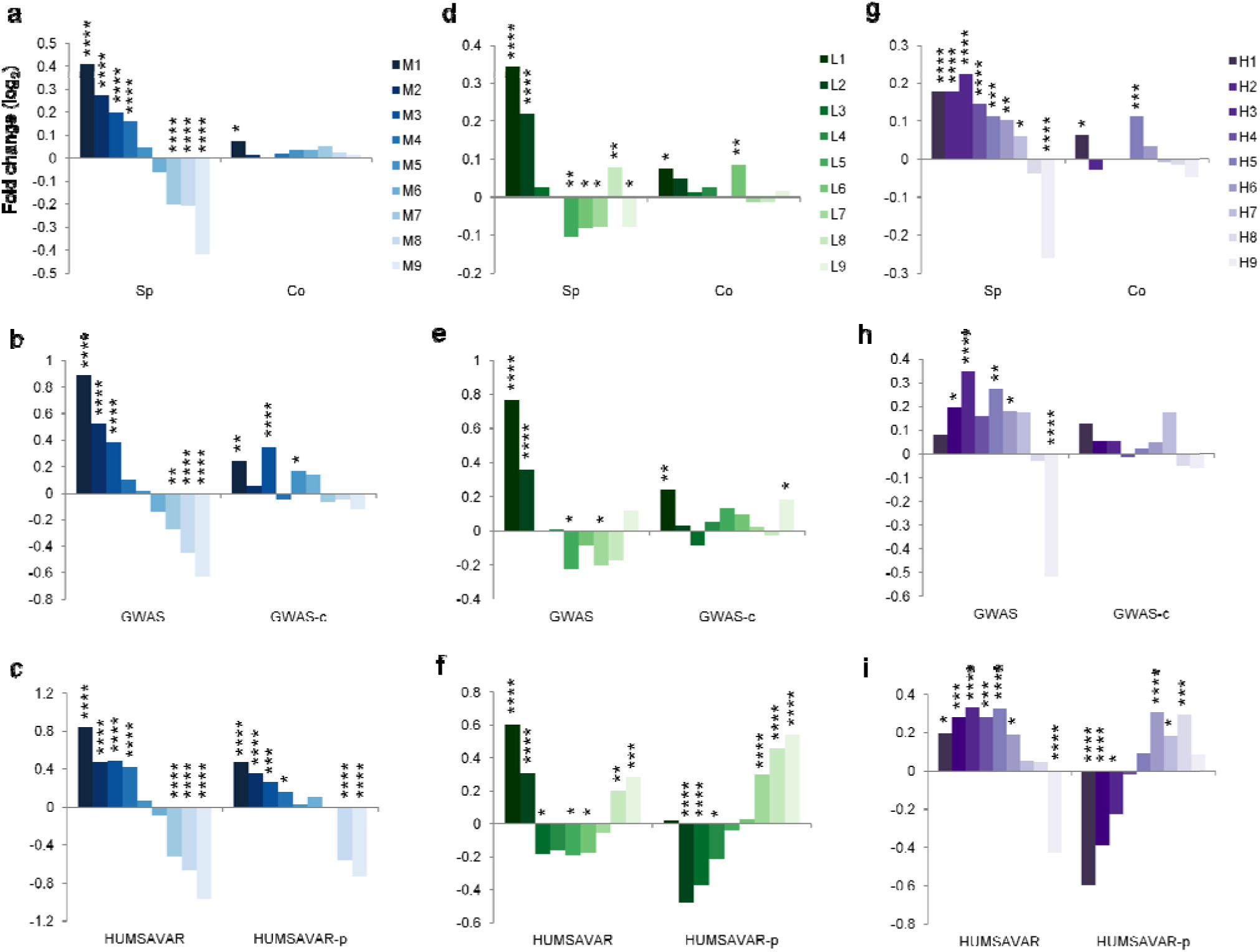
Mutational profiles of sperm and disease genes. Overlap of human sperm and disease associated genes in gene sets representing decreasing order of (**a**-**c**) mutability, M1-M9; (**b**-**d**) loss-of-function intolerance, L1-L9; and (**g**-**i**) haploinsufficiency, H1-H9. Note stronger enrichment of higher order genes in sperm (Sp) and disease (GWAS, HUMSAVAR) in general, compared to the respective controls Co, GWAS-c, and HUMSAVAR-p. Gene list sources, remarks, and overlapping genes are given in **Supplementary Table 2**. Abbreviations and other details as mentioned in the text and Fig. 1.

**Fig. 3.**
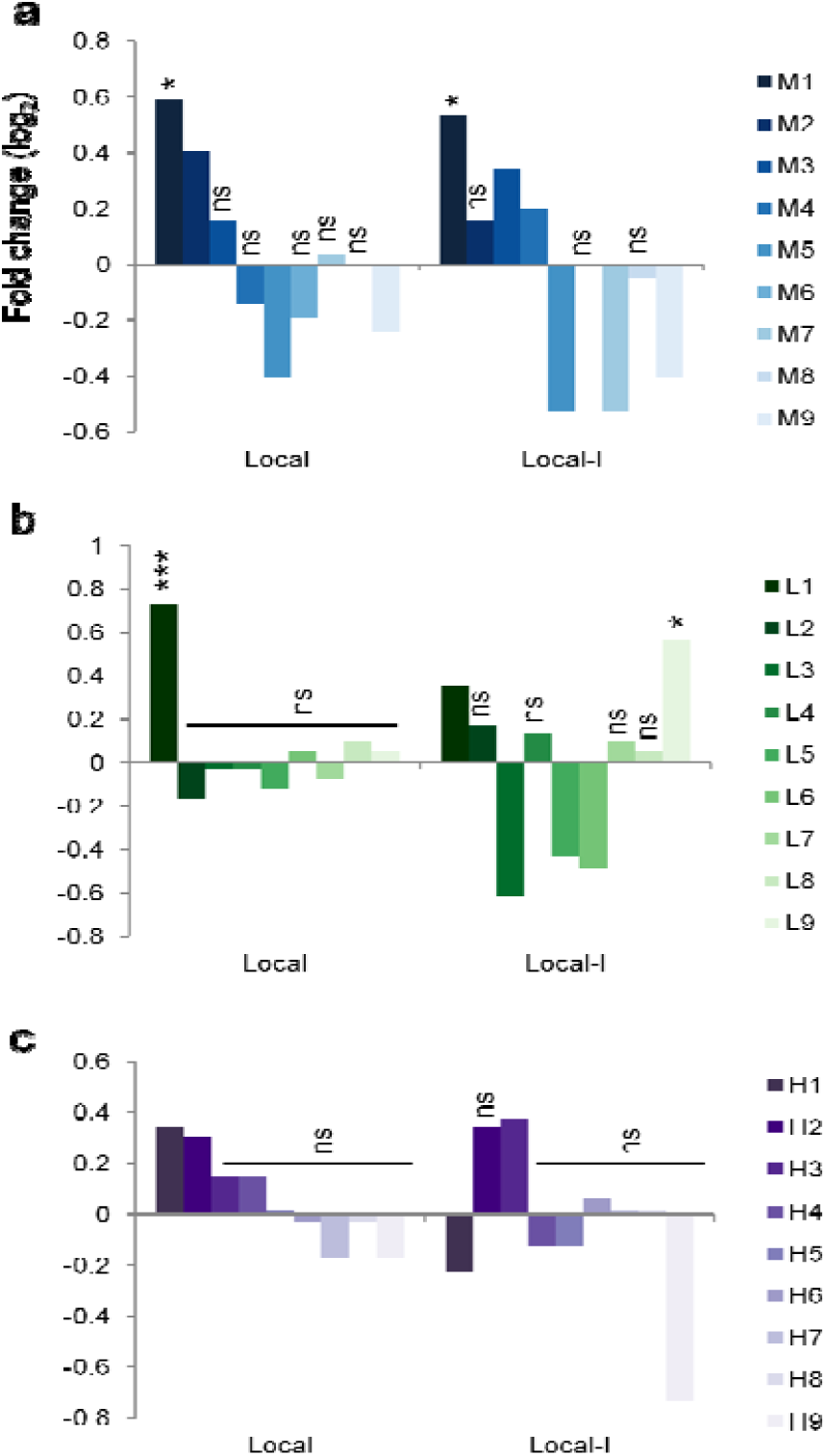
Mutational profile of adaptation associated genes. Overlap of adaptation genes in gene sets representing decreasing order of mutability (**a**), loss-of-function intolerance (**b**), and haploinsufficiency (**c**). Note greater enrichment of higher order genes in general in Local, compared to the control Local-l. Overlapping genes are listed in **Supplementary Table 3**. Abbreviations and other details as mentioned in the text, and Fig. 1 and 2.

Next, to examine mechanistic plausibility of epigenetic inheritance, I tested the null hypothesis that, in both mice and humans, the extent of overlap between combined set of differentially methylated genes in sperm and genes known to preserve sperm methylation in development is the same as that between the former and the genes in the whole genome. The alternative hypothesis was that the extent of overlap with methylation retainers is greater, supporting altered methylation state as a mediator of inheritance. For gene methylation bias in mice, a randomly selected set of database genes, matched in number with sperm genes, showing chemical induced differential methylation in somatic tissues was used as control. For human sperm analysis, the previous (**Fig. 1d, e, 2, and 3**) EWAS control was used. The developmental epigenetic gene sets were represented by demethylation resistant genes observed in blastocyst, epiblast, and primordial germ cells in mice^3,29^, genes that retain sperm methylation pattern in inner cell mass and trophectoderm cells isolated from human blastocysts, and demethylation resistance genes observed in primordial germ cells in humans^4,30^. Notably, the sperm gene set, not the control, showed overrepresentation of developmental epigenetic genes in general, in both mice (**Fig. 3a**) and humans (**Fig. 3b**), supporting the alternative hypothesis. To further examine the feasibility of paternal sperm methylome impacting offspring phenotype, I repeated the above analysis by replacing the developmental epigenetic genes with developmentally expressed genes. Again, the sperm gene set, not the control, was in general found to overrepresent genes expressed across developmental stages, in both mice (**Fig. 3c**) and humans (**Fig. 3d**). Together, these findings suggested that sperm methylome influences development and hence can mediate inheritance.

Together, the present analysis addresses key uncertainties about germline epigenetic inheritance, by supporting a role of sperm DNA methylome in development, inheritance, evolutionary adaptation, and disease. The view emerges that environmentally triggered germline epigenome changes may mediate inheritance of acquired characteristics and induce mutational events causal to genetic fixation of the traits, thereby acting as substrates for directed evolution. Providing a broad framework of environmental epigenetics, the analysis suggests that sperm DNA methylome may prove valuable in understanding genotype-phenotype correlations, identifying adaptive and disease genes, and explaining missing heritability. In general, it supports integration of epigenetic inheritance in modern evolutionary synthesis.

## Methods

Literature-wide search was conducted to manually identify relevant papers. To maintain the gene overlap analysis unbiased, only the gene lists associated with the publications or available in public databases were used, except for a single publication where SNP, not gene, list was provided. In the latter case, the host, and left- and right-flanking database genes corresponding to the given SNPs were retrieved. The gene list sources including publications and databases, along with remarks, if any, are mentioned in **Supplementary Tables 1, 2**, and **4**. MouseMine and HumanMine gene symbols were used for finding overlapping genes, with total protein-coding genes in the two databases representing whole mouse genome and whole human genome, in that order. Control gene sets were assembled using a random number generator. Hypergeometric distribution probability was used for analyzing gene overlaps, using the formula (a/b)/(c/d), where a, b, c, and d represented number of successes in the sample, sample size, number of successes in the population, population size, in that order. The population was either differentially methylated sperm genes combined or whole genome, as applicable. Bonferroni correction was applied for adjustment of overlap *p* values. Fold changes were expressed in terms of log_2_ values, with positive values indicating overrepresentation or enrichment and negative values indicating underrepresentation or depletion.

## Additional information

Supplementary Table 1. Gene list sources, remarks, and overlapping genes related to Fig. 1

Supplementary Table 2. Gene list sources, remarks, and overlapping genes related to Fig. 2

Supplementary Table 3. Overlapping genes related to Fig. 3

Supplementary Table 4. Gene list sources, remarks, and overlapping genes related to Fig. 4

**Fig. 4.**
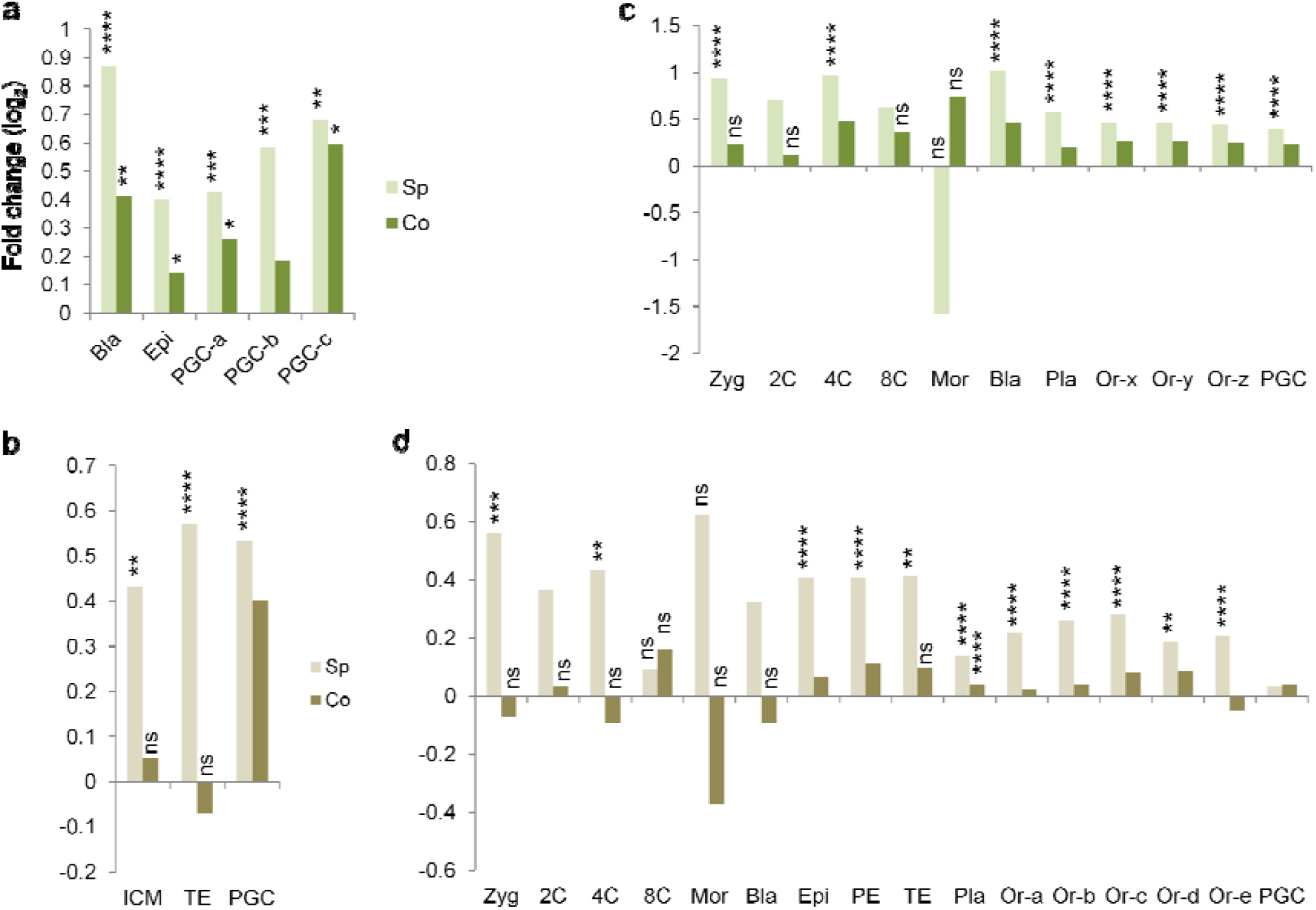
Developmental profiles of sperm genes. Fold enrichment or depletion of sperm genes in developmental epigenetic mouse (**a)** and human (**b**) genes, and in developmentally expressed mouse (**c**) and human (**d**) genes. Sp and Co (a, c) represent combined set of all 3 mouse model (LPD, CE, and H+S) genes, and differentially methylated database control genes, in that order. Bla, blastocyst; Epi, epiblast; ICM, inner cell mass; Mor, morula; Or-a, -b, -c, -d, and -e represent differential gene expression between Carnegie stages S9-S10, S10-S11, S11-S12, S12-S13, and S13-S14, in that order; Or-x, -y, and -z represent early organogenesis main cell types, main trajectories, and sub-trajectories, in that order; PE, primitive endoderm; PGC, primordial germ cells, with -a, -b, and -c indicating embryonic day 9.5, 10.5, and 11.5, in that order; Pla, placenta; TE, trophectoderm; Zyg, zygote; 2C, 4C, and 8C represent 2-cell, 4-cell, and 8-cell embryos, in that order. Note enrichment of Sp, not Co, in both epigenetic (a, b) and expressed (c, d) gene sets across species. Gene list sources, remarks, and overlapping gene sets are shown in **Supplementary Table 4**. Other abbreviations and details as mentioned in the text, and Fig. 1.

